# A single nucleotide substitution in *TaHKT1;5-D* controls shoot Na^+^ accumulation in bread wheat

**DOI:** 10.1101/2020.01.21.909887

**Authors:** Chana Borjigin, Rhiannon K. Schilling, Jayakumar Bose, Maria Hrmova, Jiaen Qiu, Stefanie Wege, Apriadi Situmorang, Chris Brien, Bettina Berger, Matthew Gilliham, Allison S. Pearson, Stuart J. Roy

## Abstract

Improving salinity tolerance in the most widely cultivated cereal, bread wheat (*Triticum aestivum* L.), is essential to increase grain yields on saline agricultural lands. A Portuguese landrace, Mocho de Espiga Branca accumulates up to 6 folds greater leaf and sheath sodium (Na^+^) than two Australian cultivars, Gladius and Scout, under salt stress. Despite high leaf and sheath Na^+^ concentrations, Mocho de Espiga Branca maintained similar salinity tolerance compared to Gladius and Scout. A naturally occurring single nucleotide substitution was identified in the gene encoding a major Na^+^ transporter TaHKT1;5-D in Mocho de Espiga Branca, which resulted in a L190P amino acid residue variation. This variant prevents Mocho de Espiga Branca from retrieving Na^+^ from the root xylem leading to a high shoot Na^+^ concentration. The identification of the tissue tolerant Mocho de Espiga Branca will accelerate the development of more elite salt tolerant bread wheat cultivars.

## Introduction

Globally, 45 million ha of irrigated and 32 million ha of dryland agricultural land is affected by salinity (FAO, 2019a). It is estimated that up to 1.2 million ha of land is lost to salinization each year (FAO, 2019a). To feed the rapidly growing human population, global food production must increase more than 70% by 2050, equating to an average increase of 44 million metric tons per year (Tester & Langridge, 2010). Bread wheat (*Triticum aestivum*) is the most widely cultivated cereal crop, in terms of area and provides one fifth of the total calories consumed worldwide (FAO, 2019b). Improving the salinity tolerance of bread wheat to maximise yields on saline agricultural land is required.

Salinity affects plants in two distinct stages. Shoot ion-independent stress (osmotic stress) arises immediately after plants are exposed to salt, inducing rapid physiological responses, such as stomatal closure and slower cell expansion, resulting in reduced plant growth (Munns & Tester, 2008). Shoot ion-dependent stress (ionic stress) has a slower onset and occurs when salt, particularly Na^+^ and Cl^−^, accumulate to high concentrations in the shoot (Munns & Tester, 2008). In this phase, reduced plant growth becomes more evident and premature leaf senescence occurs due to the toxicity of salt on cell metabolism (Munns & Tester, 2008).

Plants have three main mechanisms for tolerating salinity: osmotic stress tolerance via rapid, long distance signalling to maintain plant growth; ionic tissue tolerance by compartmentalising excessive Na^+^ or Cl^−^ in vacuoles to avoid accumulation to toxic concentrations in the cytoplasm, and the exclusion of Na^+^ from the shoot by retrieving Na^+^ from the xylem into the root or through efflux of Na^+^ into the soil to maintain a low shoot Na^+^ concentration (Munns & Tester, 2008).

The High-Affinity Potassium Transporter 1;5 (HKT1;5) is known to be responsible for retrieving Na^+^ from the transpiration stream in the root and preventing Na^+^ from accumulating to high concentrations in the shoot (Hamamoto, Horie, Hauser, Deinlein, Schroeder & Uozumi, 2015). Plant HKT proteins belong to the high-affinity K^+^/Na^+^ transporting Ktr/TrK/HKT superfamily and are divided into two groups based on a serine/glycine substitution in the first loop of the proteins (Platten, Cotsaftis, Berthomieu, Bohnert, Davenport, Fairbairn, Horie, Leigh, Lin, Luan, Maser, Pantoja, Rodriguez-Navarro, Schachtman, Schroeder, Sentenac, Uozumi, Very, Zhu, Dennis & Tester, 2006). Members of the HKT1 group with a serine residue, typically transport Na^+^, while the HKT2 group with a glycine side chain generally transport both Na^+^ and K^+^ (Platten *et al*., 2006). In Arabidopsis (*Arabidopsis thaliana*) overexpression of *AtHKT1;1* in root stellar cells reduced shoot Na^+^ accumulation by up to 64% compared to null segregants (Moller, Gilliham, Jha, Mayo, Roy, Coates, Haseloff & Tester, 2009). In bread wheat, the *AtHKT1;1* ortholog, *TaHKT1;5-D* reduces shoot Na^+^ accumulation under salinity (Byrt, Xu, Krishnan, Lightfoot, Athman, Jacobs, Watson-Haigh, Plett, Munns, Tester & Gilliham, 2014). Durum wheat (*Triticum turgidum* L.), which lacks the D-genome accumulates high concentrations of Na^+^ in the leaf (Munns, Rebetzke, Husain, James & Hare, 2003). However, when *TmHKT1;5-A* (*Nax2* locus) from a wheat relative *Triticum monococcum* L. was introduced into a commercial durum wheat, a reduction in leaf Na^+^ concentration (James, Davenport & Munns, 2006) and a 25% improvement in grain yield in the field were observed (Munns, James, Xu, Athman, Conn, Jordans, Byrt, Hare, Tyerman, Tester, Plett & Gilliham, 2012).

Breeding of salt tolerant bread wheat cultivars has focused on selecting genotypes with improved shoot Na^+^ exclusion (Ashraf & O’leary, 1996, Poustini & Siosemardeh, 2004). However, selection based on shoot Na^+^ exclusion is not always correlated with increased salinity tolerance in bread wheat (Genc, Oldach, Gogel, Wallwork, McDonald & Smith, 2013, Genc, Taylor, Lyons, Li, Cheong, Appelbee, Oldach & Sutton, 2019). Barley (*Hordeum vulgare* L.) is one of the most salt tolerant cereal crops and has much higher leaf Na^+^ concentrations than bread wheat yet is able to maintain shoot growth in saline soils (Munns & Tester, 2008, Tilbrook, Schilling, Berger, Garcia, Trittermann, Coventry, Rabie, Brien, Nguyen & Tester, 2017). The identification of a bread wheat line that accumulates high shoot Na^+^ concentrations whilst maintaining salinity tolerance, similar to barley, may accelerate the development of more salt tolerant bread wheat cultivars.

Here, we identified a Portuguese bread wheat landrace Mocho de Espiga Branca, which accumulated significantly higher leaf Na^+^ concentrations while maintaining similar salinity tolerance as current commercial elite bread wheat cultivars. A naturally occurring single nucleotide polymorphism (SNP) in *TaHKT1;5-D* prevents Mocho de Espiga Branca from retrieving Na^+^ from the root xylem, which results in a greater flux of Na^+^ to the shoot and higher accumulation of Na^+^ in leaf tissues.

## Materials and Methods

### Plant materials and growth condition

Two Australian commercial bread wheat cultivars Gladius and Scout and a set of 73 bread wheat diversity lines consisting of advanced cultivars, landraces and synthetic hexaploids were initially screened for salinity tolerance in this study (Garcia, Eckermann, Haefele, Satija, Sznajder, Timmins, Baumann, Wolters, Mather & Fleury, 2019). In the subsequent physiological characterizations, one of the 73 bread wheat diversity lines, a Portuguese landrace Mocho de Espiga Branca, and the commercial cultivars Gladius and Scout were used. Plants in all the glasshouse experiments were grown under natural light with a daytime temperature at 22 °C and 15 °C at night.

### Screening of bread wheat diversity set in a pot trial

A total of 75 lines (Gladius and Scout together with the 73 diversity lines) were screened for salinity tolerance in a fully automated conveyor system within a temperature controlled Smarthouse at the Australian Plant Phenomics Facility (The Plant Accelerator^®^, University of Adelaide, Australia; Latitude: −34.971366°, Longitude: 138.639758°) between August and September of 2014. Uniform sized seeds of each line were placed into 50 ml polypropylene tubes and imbibed in reverse osmosis (RO) water at room temperature for 4 h, tubes were drained and placed in the dark at 4 °C for three days before sowing. 3 seeds from each line were sown in a free-draining plastic pot (145 mm diameter × 190 mm height) filled with 2.5 kg of a soil mixture (50% (v/v) University of California mix, 35% (v/v) peat mix and 15% (v/v) clay loam). The pots were arranged according to a split-plot experimental design with two consecutive pots forming a main plot. The lines were allocated to the main plots and were unequally replicated with Gladius and Scout replicated 12 times, 21 randomly-selected lines of the 73 diversity lines were replicated 4 times and the remaining 52 lines were replicated three times. The subplot design randomized the two treatment levels (0 and 100 mM NaCl) in each main plot. The main plot design was a blocked row-and-column design generated using DiGGer (Coombes, 2009) and the subplot randomization was generated using dae(Brien, 2014), packages for the R statistical computing environment (R Core Team, 2014). At the emergence of the 2^nd^ leaf, plants were thinned to one uniformly sized plant per pot. At emergence of the 3^rd^ leaf, pots were loaded onto an individual cart in the Smarthouse, where they were weighed daily and automatically watered to maintain the soil water content in each pot at 17% (w/w). At the emergence of the 4^th^ leaf, 213 ml of either 0 or 170 mM NaCl solution was applied to a saucer below each pot. Plants were not watered again until the soil water content reduced below 17% (w/w), and then each pot was watered again daily to maintain a final treatment concentration of 0 or 100 mM NaCl in the respective pot.

The shoot area of each plant was non-destructively imaged using a LemnaTec 3D Scanalyzer system (LemnaTec GmbH) for a total of 15 days (4 days before and 11 days after the NaCl treatment). Three red-green-blue (RGB) images (one top view and two side view images with a 90° angle) were recorded daily for each plant to calculate the projected shoot area (PSA) in pixels. 11 days after the NaCl treatment, the 4^th^ leaf blade of each plant was collected, weighed and dried in an oven at 60 °C for two days before the dry weight was recorded. The dried 4^th^ leaf blade was subsequently used for measuring Na^+^, K^+^ and Cl^−^ concentrations.

### Determining leaf blade, leaf sheath and root Na^+^ concentration in hydroponics

To measure the Na^+^, K^+^ and Cl^−^ concentrations in the 4^th^ leaf blade and sheath and roots, Mocho de Espiga Branca, Gladius and Scout were grown using a fully supported hydroponics system under three concentrations of NaCl (0, 150 and 200 mM) in a controlled glasshouse at The Plant Accelerator^®^ between June and August 2017. The hydroponic system was equipped with a trolley fitted with two growth trays each containing 42 tubes filled with polycarbonate pellets and a tank containing 80 l of a modified Hoagland solution (mM): NH_4_NO_3_ (0.2); KNO_3_ (5.0); Ca(NO_3_)_2_·4H_2_O (2.0); MgSO_4_·7H_2_O (2.0); KH_2_PO_4_ (0.1); Na_2_Si_3_O_7_ (0.5); NaFe(III)EDTA (0.05); MnCl_2_·4H_2_O (0.005); ZnSO_4_·7H_2_O (0.01); CuSO_4_·5H_2_O (0.0005) and Na_2_MoO_3_·2H_2_O (0.0001). Uniform sized seeds from each genotype were surface sterilized using ultraviolet (UV) light for two min and germinated in petri dishes on moist filter paper for 4 days at room temperature before transplanting. 14 replicates from each cultivar were grown in each treatment trolley. The NaCl treatment was applied at the emergence of the 4^th^ leaf by adding 116.88 g of NaCl twice daily for a 25 mM NaCl increment until a final concentration of either 150 mM or 200 mM was reached, and no NaCl was added to the 0 mM NaCl treatment. 3.8 g of supplemental CaCl_2_·2H_2_O was added into the 150 and 200 mM NaCl treatment tanks at each 25 mM NaCl increment in order to maintain constant Ca^2+^ activity in all three treatments. The plants were irrigated by the nutrient solution in a 20 min flood and rain cycle and the complete nutrient solution was replaced every seven days. The pH of the solution was maintained between 6.5 and 7.0 throughout the experiment using a 3.2% (v/v) HCl solution and a portable waterproof specific Ion–pH–mV–Temperature meter (Modal WP-90, TPS Pty Ltd, Australia). After 21 days of NaCl treatment, the 4^th^ leaf blade and sheath and the remaining shoots were sampled separately and weighed. Plant roots were weighed after sampling and approximately 5 cm of the root tip was used for RNA extraction. The roots from the 150 mM and 200 mM treatments were rinsed in 10 mM CaSO_4_ solution and blotted on paper towel to remove traces of NaCl before sampling. The weighed 4^th^ leaf blade, 4^th^ leaf sheath, shoots and roots were dried in an oven at 60 °C for two days to record the dry weight. The dried 4^th^ leaf blade and sheath and root tissue were used for the subsequent Na^+^, K^+^ and Cl^−^ concentration analysis.

### Na^+^, K^+^ and Cl^−^ concentration analysis

The harvested dried 4^th^ leaf blade and root samples were digested in 10 ml of 1% (v/v) HNO_3_ (v/v), and the 4^th^ leaf sheath was digested in 5 ml of 1% (v/v) HNO_3_ (v/v) at 85 °C for 4 h in a SC154 HotBlock® (Environmental Express Inc., South Carolina, US). Na^+^ and K^+^ concentrations were measured using a flame photometer (Model 420; Sherwood Scientific Ltd., Cambridge, UK), and Cl^−^ concentration was measured using a chloride analyzer (Model 926; Sherwood Scientific Ltd., Cambridge, UK).

### RNA extraction and cDNA synthesis

The harvested 5 cm root tips from the hydroponics experiment was snapped frozen in liquid nitrogen and stored at −80 °C until RNA extraction. The root tissue was ground to a fine powder using a 2010 Geno/Grinder® (SPEX SamplePrep) at 1000 ×g for 30 sec, and total RNA was extracted from the tissue powder using a Direct-Zol RNA MiniPrep kit with DNase I treatment (Zymo Research) according to the manufacturer’s instruction. Final elution was performed with 40 µl DNA/RNAase-Free water supplied and the eluted RNA was subsequently quantified using a ND-1000 Spectrophotometer (NanoDrop Technologies) and quantified on a 1% (w/v) agarose gel (Bioline) by electrophoresis. cDNA synthesis was performed on 500 ng of RNA using High Capacity cDNA Reverse Transcription Kit (Thermo Fisher Scientific) according to the manufacturer’s instruction in a 20 µl reaction and stored at −20 °C until use.

### *TaHKT1;5-D* coding sequence amplification and sequencing

The entire coding sequence of the TaHKT1;5-D gene from Mocho de Espiga Branca, Gladius and Scout were amplified using Phusion® High-Fidelity DNA polymerase (New England Biolabs) following the manufacturer’s instruction. The primers used for PCR amplification were: forward primer *cTaHKT1;5-D_FP_1* (5’-ATGGGTTCTTTGCATGTCTCCT-3’) and reverse primer *cTaHKT1;5-D_RP_1551* (5’-TTATACTATCCTCCATGCCTCGC-3’) (Table S3). PCR was conducted on a T100™ Thermal Cycler (Bio-Rad) using the following conditions: initial denaturation at 98 °C for 30 s, 35 cycles of 98 °C for 30 s, 64 °C for 30 s, 72 °C for 1 min, final extension at 72 °C for 10 min and held at 4 °C. The PCR product was visualized on a 1% (w/v) agarose gel (Bioline) by electrophoresis at 90 V for 1 h and the 1.5 kb target band was collected for purification using NucleoSpin® Gel and PCR Clean-up kit (Macherey-Nagel) according to the manufacturer’s instruction prior to sequencing. Three replicates from each of Mocho de Espiga Branca, Gladius and Scout TaHKT1;5-D coding sequence were tested with primers listed in Table S3 for Sanger sequencing carried out at the Australian Genome Research Facility (AGRF, South Australia).

### *TaHKT1;5-D* gene expression in roots

To determine *TaHKT1;5-D* expression in Mocho de Espiga Branca, Gladius and Scout, a PCR was conducted on the synthesised cDNA in a 25 µl reaction consisting of 1 µl cDNA, 0.5 µl each of 10 µM forward primer 5’-CGACCAGAAAAGGATAACAAGCAT-3’ and reverse primer 5’-AGCCAGCTTCCCTTGCCAA-3’, 5 µl *Taq* 5× Master Mix (New England Biolabs) and 18 µl Milli-Q water (18.2 MΩ cm). The final PCR products were visualized on a 1% (w/v) agarose gel (Bioline) by electrophoresis at 90 V for 1 h. The targeted *TaHKT1;5-D* product was 283 bp. The *TaGAP* gene (230 bp) was used as a positive control and a Milli-Q water sample was included as a negative control.

To quantify *TaHKT1;5-D* expression in the root tissue of Mocho de Espiga Branca, Gladius and Scout, Real-time PCR was performed on five replicates from each of the cultivars using an Applied Biosystems™ QuantStudio™ 6 Flex (Life Technologies). The reaction was performed in a 10 µl reaction consisting of two µL 1:20 diluted synthesised cDNA, 0.5 µl each of 10 µM forward and reverse primers stated above, 5 µl KAPA SYBR FAST 2× Master Mix (Sigma-Aldrich) and 2 µl Milli-Q water. The 283 bp final product from each cultivar was confirmed by Sanger sequencing carried out at AGRF (South Australia).

### DNA extraction and quantification

Genomic DNA (gDNA) extraction of Mocho de Espiga Branca, Gladius, Scout and 70 bread wheat diversity lines was performed using a phenol/chloroform/iso-amyl alcohol extraction method as described elsewhere with modifications (Rogowsky, Guidet, Langridge, Shepherd & Koebner, 1991). Briefly, the leaf tissue was frozen in a 10 ml tube at −80 °C and ground to a fine powder using a 2600 Cryo-Station (SPEX^®^ SamplePrep) and a 2010 Geno/Grinder (SPEX^®^ SamplePrep) at 1000 ×g for 30 s. 2 ml of gDNA extraction buffer [1% (w/v) sarkosyl, 100 mM Tris-HCl, 100 mM NaCl, 10mM EDTA, 2% (w/v) insoluble Polyvinylpolypyrrolidone] was added to the ground tissue, vortexed, followed by the addition of 2 ml phenol/chloroform/iso-amyl alcohol (25:24:1). The sample was placed on ice for 20 min and vortexed thoroughly in every 5 min before centrifuging at 3630 ×g for 15 min. The supernatant was transferred into a labelled BD Vacutainer™ SST™ II Advance tube (Becton, Dickinson and Company, New Jersey, US) and 2 ml of phenol/chloroform/iso-amyl alcohol (25:24:1) was added. The sample was vortexed and centrifuged as above, and the supernatant was collected into a new 10 ml tube. The gDNA was precipitated using 2 ml of 100% (v/v) iso-propanol and 200 µl of 3 M sodium acetate (pH 4.8) and washed using 1 ml of 70% (v/v) ethanol before re-suspending overnight in 80 µl of R40 at 4°C. The re-suspended gDNA was quantified using a ND-1000 Spectrophotometer (NanoDrop Technologies).

### Cleaved Amplified Polymorphic Sequence (CAPS) assay and genotyping

A CAPS marker tsl2SALTY-4D was designed to confirm the allele effect of the SNP (T/C) identified in Mocho de Espiga Branca *TaHKT1;5-D* for high leaf Na^+^ concentration in comparison to Gladius and Scout. The extracted DNA from Mocho de Espiga Branca, Gladius, Scout and 68 bread wheat diversity lines were used for a PCR to amplify a 945 bp DNA fragment containing SNP. The PCR analysis was conducted in a 10 µl reaction consisting of 1 µg of DNA, 0.24 µl each of 10 µM forward primer 5’**-**ATGGGTTCTTTGCATGTCTCCT**-**3’ and reverse primer 5’**-**CGCTAGCACGAACGCCG**-**3’, 2 µl of *Taq* 5× Master Mix (New England Biolabs) and Milli-Q water. The reaction was performed on a T100™ Thermal Cycler (Bio-Rad) using the following conditions: initial denaturation at 95 °C for 4 min, 35 cycles of 95 °C for 30 s, 56 °C for 30 s, 68 °C for 1 min, final extension at 68 °C for 5 min and held at 12 °C. The PCR amplification was followed by digestion using the restriction enzyme *FauI*. It was conducted in a 10 µl reaction consisting of 1 µg of the PCR products, 0.4 µl of *FauI* enzyme (New England Biolabs), 1 µl of 10×CutSmart^®^ Buffer (New England Biolabs) and Milli-Q water. The digestion was performed on a T100™ Thermal Cycler (Bio-Rad) for 1 h at 55 °C followed by 20 min of inactivation at 65 °C and held at 12 °C. The digested product was visualized on a 2% (w/v) agarose gel (Bioline) by electrophoresis at 90 V for 90 min and the genotype of each line at the SNP position was confirmed according to the product bands on the gel. Lines which had the C:C genotype had two fragments present at 573 and 372 bp, whilst lines carrying the T:T genotype had a single fragment present.

### Construction of 3D molecular models of TaHKT1;5-D and TaHKT1;5-D L190P

The most suitable template for wheat HKT1;5 transporter proteins was the *B. subtilis* KtrB K^+^ transporter (Protein Data Bank accession 4J7C, chain I) (Vieira-Pires, Szollosi & Morais-Cabral, 2013) as previously identified (Xu, Waters, Byrt, Plett, Tyerman, Tester, Munns, Hrmova & Gilliham, 2018). The K^+^ ion in KtrB was substituted by Na^+^ during modelling of TaHKT1;5 proteins. 3D models of TaHKT1;5-D and TaHKT1;5-D L190P in complex with Na^+^ were generated with Modeller 9v19 (Sali & Blundell, 1993) as described previously (Cotsaftis, Plett, Shirley, Tester & Hrmova, 2012, Waters, Gilliham & Hrmova, 2013) incorporating Na^+^ ionic radii (Xu *et al*., 2018) taken from the CHARMM force field (Brooks, Brooks, Mackerell, Nilsson, Petrella, Roux, Won, Archontis, Bartels, Boresch, Caflisch, Caves, Cui, Dinner, Feig, Fischer, Gao, Hodoscek, Im, Kuczera, Lazaridis, Ma, Ovchinnikov, Paci, Pastor, Post, Pu, Schaefer, Tidor, Venable, Woodcock, Wu, Yang, York & Karplus, 2009), on the Linux station running the Ubuntu 12.04 operating system. Best scoring models (from an ensemble of 50) were selected based on the combination of Modeller Objective (Shen & Sali, 2006) and Discrete Optimised Protein Energy (Eswar, Eramian, Webb, Shen & Sali, 2008) functions, PROCHECK (Laskowski, Macarthur, Moss & Thornton, 1993), ProSa 2003(Sippl, 1993) and FoldX (Schymkowitz, Rousseau, Martins, Ferkinghoff-Borg, Stricher & Serrano, 2005). Structural images were generated in the PyMOL Molecular Graphics System V1.8.2.0 (Schrődinger LLC, Portland, OR, USA). Calculations of angles between selected α-helices in HKT1;5 models were executed in Chimera (Pettersen, Goddard, Huang, Couch, Greenblatt, Meng & Ferrin, 2004) and evaluations of differences (ΔΔG = ΔGmut-ΔGwt) of Gibbs free energies was performed with FoldX (Schymkowitz *et al*., 2005). Sequence conservation patterns were analysed with ConSurf (Celniker, Nimrod, Ashkenazy, Glaser, Martz, Mayrose, Pupko & Ben-Tal, 2013, Landau, Mayrose, Rosenberg, Glaser, Martz, Pupko & Ben-Tal, 2005) based on 3D models of TaHKT1;5-D using 370 sequences at the sequence identities of 30% and higher (specifications: HMMMER homolog search algorithm, UNIREF-90 Protein database with the E-value cut-off of 1×10^−4^, Bayesian Model of substitution for proteins.

### Characterization of TaHKT1;5-D from Mocho de Espiga Branca and Gladius in *X. laevis* oocytes

Na^+^ transport properties of TaHKT1;5-D from Mocho de Espiga Branca and Gladius were characterized in *X. laevis* oocytes using two-electrode voltage clamping (TEVC) as previously described (Byrt *et al*., 2014, Munns *et al*., 2012). pGEMHE-DEST containing *TaHKT1;5-D* was linearized using sbfI-HF (New England Biolabs) before synthesising cRNA using the mMESSAGE mMACHINE T7 Kit (Ambion) following manufacturer’s instructions. 46 nl/23 ng of the cRNA from Mocho de Espiga Branca or Gladius, or equal volumes of RNA-free water (H_2_O control) were injected into oocytes. Injected oocytes were incubated for 48 h and TEVC was carried out as described (Munns *et al*., 2012). Membrane currents were recorded in the HMg solution (6 mM MgCl_2_, 1.8 mM CaCl_2_, 10 mM MES, and pH 6.5 adjusted with a Tris base) ± Na^+^ glutamate. The osmolality of the solution was adjusted to 240 - 260 mOsmol/Kg using mannitol and a micro-osmometer (Model 210, Fiske Associates Inc, USA).

### Subcellular localisation of TaHKT1;5-D

Transient expression of fluorescent fusion proteins was performed as previously described (Henderson, Wege, Qiu, Blackmore, Walker, Tyerman, Walker & Gilliham, 2015). In brief, *TaHKT1;5-D* coding sequences of Mocho de Espiga Branca and Gladius were recombined into pMDC43 (Curtis & Grossniklaus, 2003) to generate N-terminally green fluorescent protein (GFP) tagged proteins. The red fluorescent protein (RFP) tagged plasma membrane marker nCBL1-RFP that was used for co-localisation (Batistic, Sorek, Schultke, Yalovsky & Kudla, 2008). All constructs were transformed into *Agrobacterium tumefaciens* strain Agl-1 and agro-infiltration was performed on fully expanded leaves of 4 to 6 weeks old tobacco (*Nicotiana benthamiana*) plants (Henderson *et al*., 2015). After two to three days, leaf sections were imaged using a Nikon A1R Confocal Laser-Scanning Microscope equipped with a 63× water objective lens and NIS-Elements C software (Nikon Corporation). Excitation/emission conditions were GFP (488 nm/500–550 nm) and RFP (561 nm/570–620 nm).

### Xylem sap Na^+^ and K^+^ concentration analysis

Xylem sap was collected from hydroponically grown plants of Mocho de Espiga Branca and Gladius at 0 and 150 mM NaCl after 21 days from the start of the NaCl treatment. The shoot was cut off at the base of the plant and inserted into a Scholander-type pressure chamber (Model 1005, PMS Instrument Company, USA) to extrude the xylem sap by slowly filling the chamber with compressed air. The sap was immediately collected into a clean, pre-weighed 1.5 ml tube. From the 0 mM NaCl treatment, xylem sap of seven plants from each cultivars was collected, and xylem sap of eight Mocho de Espiga Branca and seven Gladius plants was collected from the 150 mM NaCl treatment. Tubes containing the xylem sap was weighed for each plant and the samples were stored at 4°C until Na^+^ and K^+^ concentrations were measured using a flame photometer (Model 420, Sherwood Scientific Ltd., Cambridge, UK).

### Net and total Na^+^, K^+^ and H^+^ flux analyses using Microelectrode Ion Flux Estimation (MIFE)

To investigate whether there were differences in Na^+^, K^+^ and H^+^ transport in the plant roots between Mocho de Espiga Branca and Gladius, net fluxes of Na^+^, K^+^ and H^+^ were measured at root maturation and elongation zones using the non-invasive MIFE technique (University of Tasmania, Hobart, Australia) (Bose, Rodrigo-Moreno, Lai, Xie, Shen & Shabala, 2015, Newman, 2001).

Root Na^+^ retrieval measurements were performed at the elongation zone (approximately 600 µm from the root cap). Uniform sized sterilized seeds from Mocho de Espiga Branca and Gladius were germinated on moist filter paper in Petri dishes covered with aluminium foil, at 4 °C overnight and then placed at room temperature in the dark for three days before transplanting. 12 seedlings from each cultivar were transplanted into a hydroponic tank containing 10 L modified Hoagland solution as previously described. NaCl was added into the solution at an increment of 25 mM (14.16 g) twice daily for two days to achieve a final concentration of 100 mM NaCl. The pH of the solution was maintained between 6.5 and 7.0 throughout the experiment using 3.2% (v/v) of the HCl solution and a portable waterproof specific Ion–pH–mV–Temperature meter (Modal WP-90, TPS Pty Ltd, Australia). Two days after being exposed to 100 mM NaCl, the entire roots of Mocho de Espiga Branca (*n* = 9) and Gladius (*n* = 8) were first preconditioned in BSM solution containing 100 mM NaCl (0.2 mM KCl + 0.1 mM CaCl_2_·2H_2_O + 100 mM NaCl) in a Petri dish for 30 min and then the primary root was immobilized on a 10 ml perspex measuring chamber containing 7 ml of the same BSM medium. The steady-state fluxes were measured for 5 min in the initial BSM solution before changing the bathing solution to 7 ml of the new BSM medium solution containing 0.6 mM NaCl (0.2 mM KCl + 0.1 mM CaCl_2_·2H_2_O + 0.6 mM NaCl). The resulting fluxes were measured for 25 min and the integral of each replication was added to derive the cumulative total fluxes. The osmolality of the two BSM solutions and the Hoagland solution were maintained between 208 and 222 mOsmol/kg using mannitol and a vapour pressure osmometer (Model 5500, Wescor, Inc., USA).

Root Na^+^ uptake measurements were performed at the maturation zone (beyond 2.5 cm from root tip), uniform sized sterilized seeds were germinated as described above, placed at 4 °C overnight and placed at room temperature in the dark for three days before measurement. The primary root of Mocho de Espiga Branca (*n* = 12) and Gladius (*n* = 11) seedlings was immobilized on a 10 ml perspex measuring chamber containing 6 ml of the BSM solution (0.2 mM KCl + 0.1 mM CaCl_2_·2H_2_O + 0.6 mM NaCl) and pre-conditioned for 20 min before recording steady-state Na^+^, K^+^ and H^+^ fluxes for 5 min then 100 mM NaCl was added and the resulting fluxes were recorded for 25 min.

### Statistical analysis

Prism 7 for Windows (version 7.02; GraphPad Software, Inc.) was used to generate graphs. GenStat^®^ 15^th^ edition for Microsoft Windows (version 15.3.09425; VSN International Ltd, UK) was used to perform an Analysis of Variance (ANOVA) and Tukey’s multiple comparison test was used to determine which means were significantly different at a probability level of *p* ≤ 0.05.

## Results

### Mocho de Espiga Branca has high leaf and sheath Na^+^ concentrations

Screening of 75 bread wheat accessions for salinity tolerance under 100 mM NaCl, identified a Portuguese landrace, Mocho de Espiga Branca, which had a similar salinity tolerance but accumulated 7 folds higher Na^+^ concentration in the 4^th^ leaf compared to all other lines, including Gladius and Scout (Figure 1 and Table S1). The 4^th^ leaf K^+^ and Cl^−^ concentrations of the 75 wheat lines were comparable under 100 mM NaCl (Table S1).

**Figure 1.**
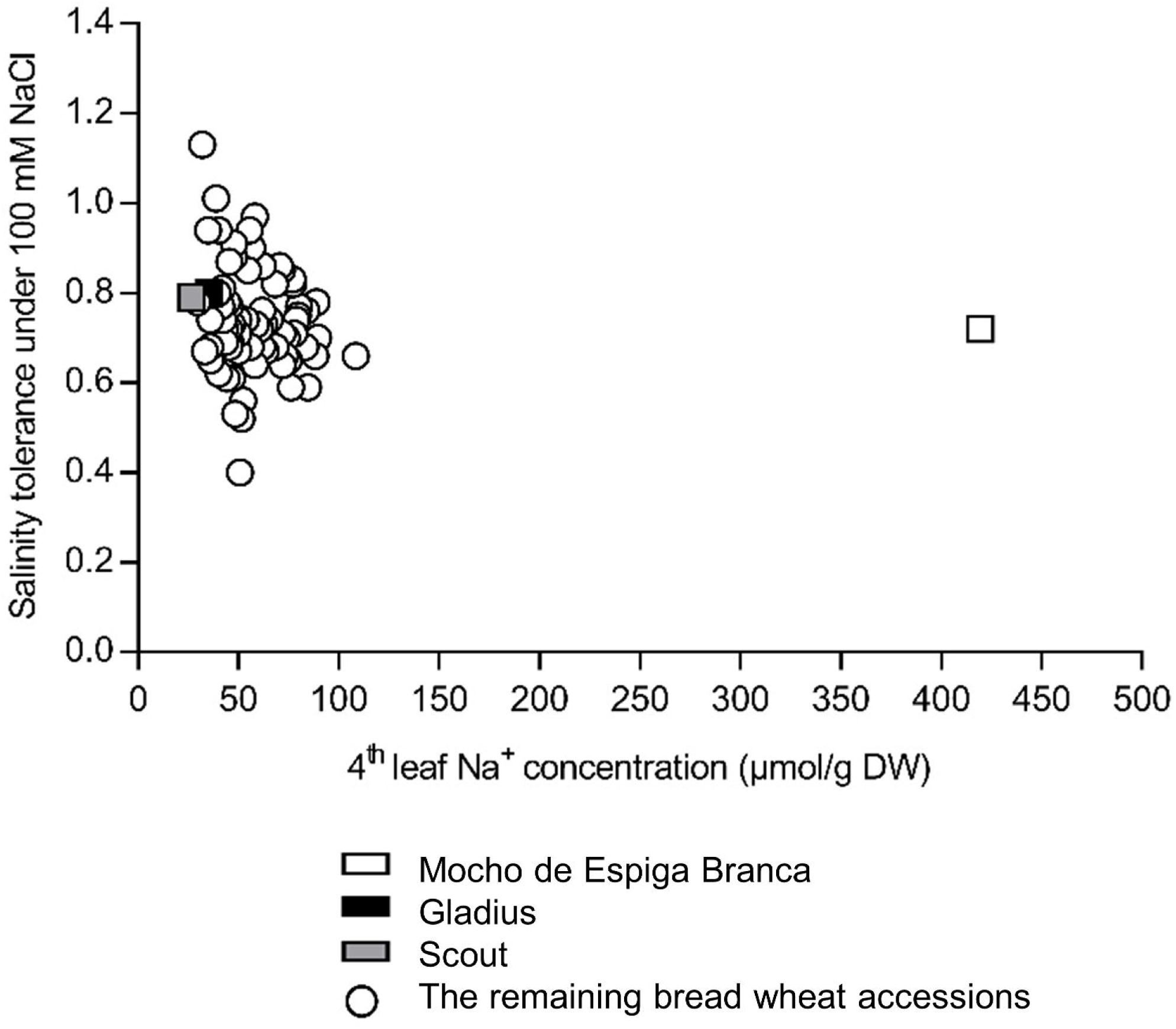
Salinity tolerance and 4^th^ leaf Na^+^ concentration of Mocho de Espiga Branca relative to 72 bread wheat diversity lines and two Australian cultivars Gladius and Scout in soil with 100 mM NaCl. The 4^th^ leaf Na^+^ concentration is determined 11 days after treatment with 100 mM NaCl. Salinity tolerance is defined as projected shoot area (PSA) under 100 mM NaCl relative to 0 mM NaCl determined from the final day of imaging. Data presented as means (*n* = 3-4 except for Gladius and Scout, where *n* = 12). The standard error of the mean (SEM) for the 4^th^ leaf Na^+^ concentration is presented in Table S1.

In hydroponics, Mocho de Espiga Branca maintained a similar shoot and root biomass to Gladius and Scout at 0, 150 and 200 mM NaCl (Figure 2a and Figure S1a,b), and all three cultivars had comparable salinity tolerance at 150 mM (0.69, 0.64 and 0.60, respectively) and 200 mM NaCl (0.39, 0.49 and 0.39, respectively) (Figure S1c,d). The 4^th^ leaf blade and sheath Na^+^ concentrations were up to 6 folds higher in Mocho de Espiga Branca than Gladius and Scout at 150 mM NaCl (Figure 2b,c). At 200 mM NaCl, 4^th^ leaf blade and sheath Na^+^ concentrations in Mocho de Espiga Branca were 5 folds greater compared to Gladius and Scout (Figure 2b,c). There was no difference in root Na^+^ concentration at all NaCl treatments (Figure 2d).

**Figure 2.**
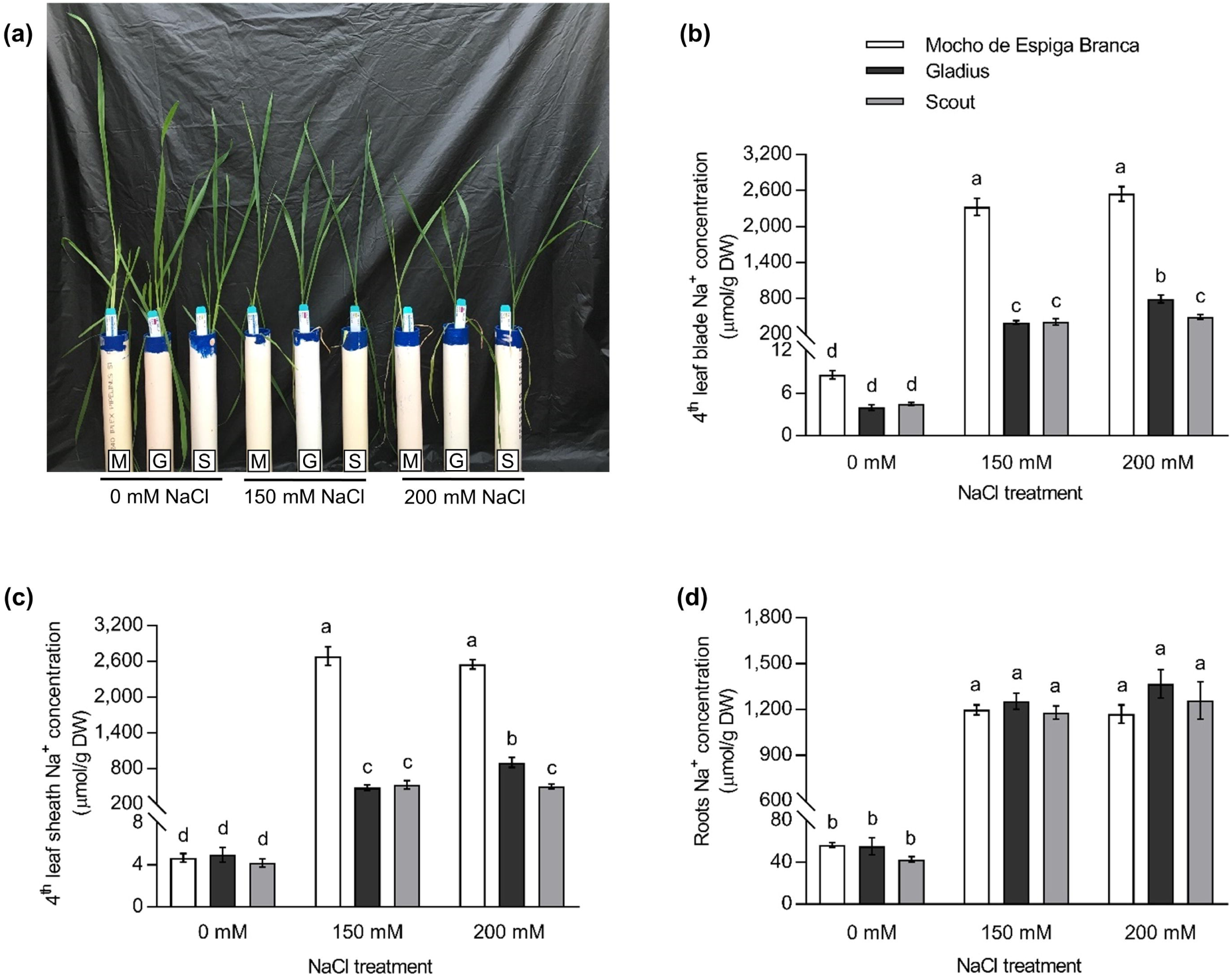
Na^+^ concentration in the 4^th^ leaf blade, sheath, and roots of Mocho de Espiga Branca, Gladius and Scout in hydroponics. **(a)** Representative image of 6 weeks old plants from the hydroponic experiment with 0, 150 and 200 mM NaCl treatments applied at the emergence of the 4^th^ leaf. M = Mocho de Espiga Branca, G = Gladius and S = Scout. Na^+^ concentration in **(b)** 4^th^ leaf blade; **(c)** 4^th^ leaf sheath and **(d)** roots determined 21 days after treatments with 0, 150 and 200 mM NaCl. Data presented as means ± SEM (*n* = 14). Bars with different letters indicate significant differences determined by two-way ANOVA with Tukey’s multiple comparison test at *p* ≤ 0.05.

All three cultivars had similar K^+^ concentrations in the 4^th^ leaf blade and sheath under 0 mM NaCl (Figure S2a). At 150 and 200 mM NaCl, Mocho de Espiga Branca accumulated 70-79% less K^+^ in the blade and 61-67% less K^+^ in the sheath compared to Gladius and Scout (Figure S2a, b). The root K^+^ concentration in Mocho de Espiga Branca was similar to Scout but significantly lower than Gladius under 0 mM NaCl, while no differences were observed between the three cultivars under 150 and 200 mM NaCl (Figure S2c).

The 4^th^ leaf blade and sheath Cl^−^ concentrations in Mocho de Espiga Branca, Gladius and Scout were similar under 0 mM NaCl (Figure S2d,e). Mocho de Espiga Branca accumulated significantly higher Cl^−^ than Gladius and Scout in both tissues under 150 and 200 mM NaCl (Figure S2d,e). There were no significant differences in root Cl^−^ concentrations at 0 mM NaCl (Figure S2f). Mocho de Espiga Branca had significantly higher Cl^−^ than Gladius and Scout at 150 mM NaCl but only significantly higher than Scout at 200 mM NaCl (Figure S2f).

### A natural single nucleotide substitution in the *TaHKT1;5-D* gene of Mocho de Espiga Branca alters Na^+^ transport properties of the protein

A natural single SNP (T/C) in the coding sequence of *TaHKT1;5-D* was identified at the 569^th^ base pair in Mocho de Espiga Branca, while the sequence of Gladius and Scout was identical to Chinese Spring (Figure 3a). The SNP in Mocho de Espiga Branca resulted in an amino acid residue change from Leucine (L) to Proline (P) at the 190^th^ residue in the Na^+^ transporter protein TaHKT1;5-D (Figure 3a). This L190P variant residue is predicted to be located on the 4^th^ transmembrane α-helix in the area of the second glycine residue of the S78-G233-G353-G457 selectivity filter motif (Figure 3b). Expression analysis of the *TaHKT1;5-D* in Mocho de Espiga Branca, Gladius and Scout showed no significant difference in expression (Figure S3a,b). A cleaved amplified polymorphic sequence (CAPS) marker tsl2SALTY-4D designed to this SNP in *TaHKT1;5-D* was used to genotype 71 diversity lines (Figure S3c and Table S2). Mocho de Espiga Branca carried the C:C allele responsible for the TaHKT1;5-D L190P variation, while all other lines had the T:T allele as Gladius and Scout (Figure S3c and Table S2).

**Figure 3.**
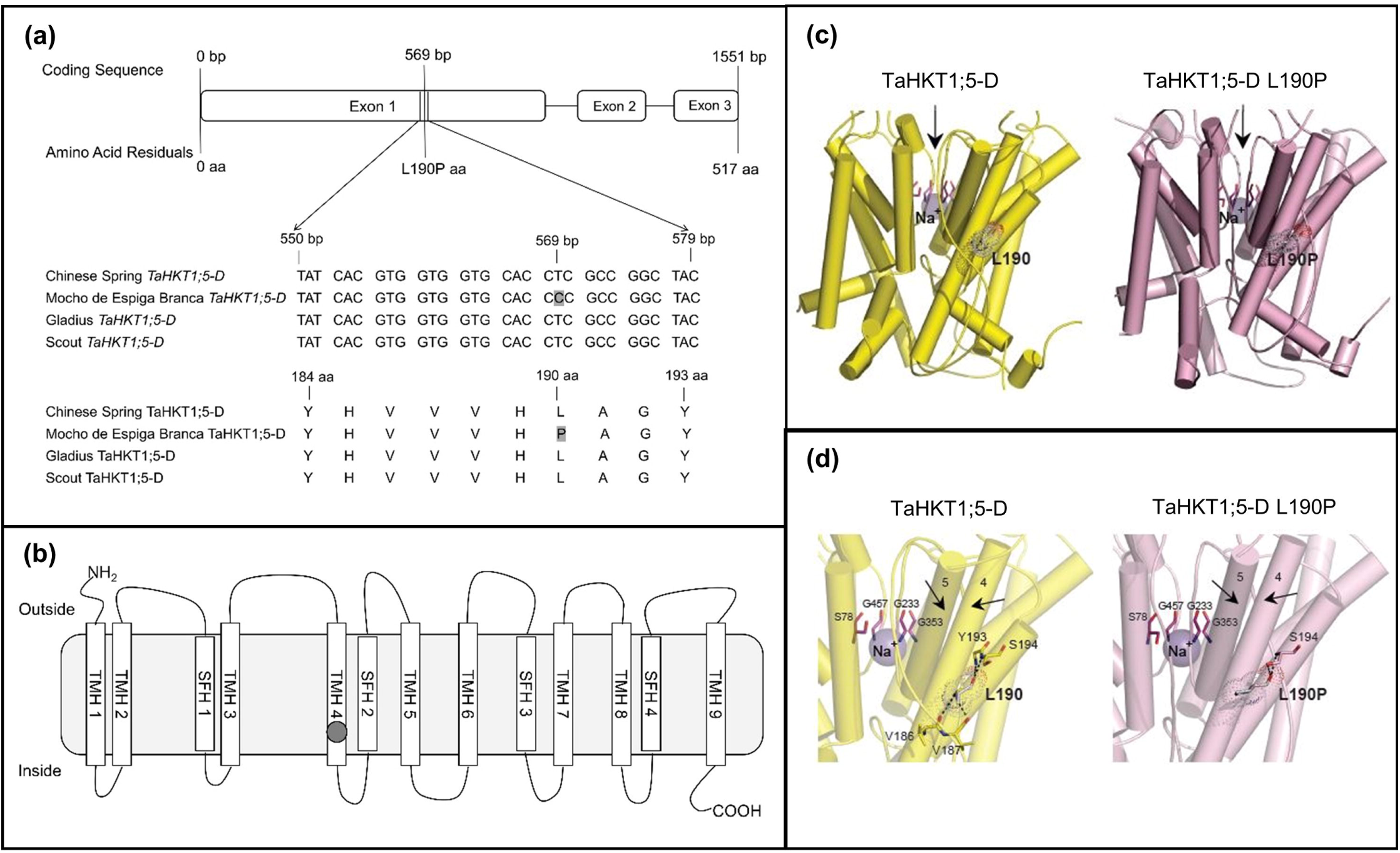
A SNP in Mocho de Espiga Branca *TaHKT1;5-D* results in a L190P amino acid residue variation in the Na^+^ transporter TaHKT1;5-D. **(a)** Partial alignment of *TaHKT1;5-D* coding and amino acid sequences in Mocho de Espiga Branca, Gladius and Scout compared to Chinese Spring. **(b)** Schematic of the TaHKT1;5-D protein showing the transmembrane α-helices (TMH 1-9) and selectivity filter α-helices (SFH 1-4) adapted from Xu et al. (2018); the Na^+^ selectivity filter motif S78-G233-G353-G457 indicated in blank circles and location of the L190P variant indicated in a grey circle. **(c)** Molecular models of TaHKT1;5-D (left, yellow) and TaHKT1;5-D L190P (right, salmon) transport proteins in cartoon representations with cylindrical α-helices illustrating 3D folds. Constrictions in selectivity filters are bound by four residues (cpk magenta sticks) that contain Na^+^ ions (violet spheres). Black arrows illustrate directional flows of Na^+^ that are likely to enter the permeation trajectory by-passing selectivity filter constrictions. Variant residues L190 and L190P (cpk sticks and dots, bold types) between wild-type TaHKT1;5-D and the L190P mutant are indicated; the dots illustrate volumes of van der Waals radii. **(d)** Detailed views of α-helices, which neighbour selectivity filter constriction, containing Na^+^ (violet spheres), located near selectivity filter residues S78, G233, G353, G457 (cpk magenta) for TaHKT1;5-D (left) and the L190P mutant (right), which are crucial for permeation function. In each protein, polar contacts (cpk sticks and dots) of L190 (TaHKT1;5-D) and L190P (TaHKT1;5-D L190P) positioned on α-helices 4, are indicated by dashed lines (separations between 2.6 Å and 3.2 Å).

3D molecular modelling revealed that overall folds of TaHKT1;5-D and TaHKT1;5-D L190P were similar (Figure 3c,d). Detailed analysis of the microenvironments around α-helix 4 and α-helix 5 (two black arrows pointing to each other in Figure 3d), revealed that L190 in α-helix 4 of TaHKT1;5-D (Figure 3d left) established four polar contacts at separations between 2.6 Å to 3.2 Å with V186, V187, Y193 and S194 neighbouring residues. These extensive polar contacts were not formed in the TaHKT1;5-D L190P variant (Figure 3d right), which only established one polar contact at the separation at 2.7 Å. In TaHKT1;5-D, the packing angle between α-helix 4 (carrying L190P) and the neighbouring α-helix 5 was 16° sharper than that in TaHKT1;5D L190P (Figure 3d). Sequence conservation patterns, based on 3D models of TaHKT1;5-D revealed that the P190 position in TaHKT1;5-D could not be found in databases, meaning that L190 could only be replaced by F, G, L, V, I, M, A, K and T, but not by P. Evaluation of differences of Gibbs free energies of TaHKT1;5-D revealed that the L190P mutation was energetically highly unfavourable (highly destabilising), and that the reverse mutation (P190 into L190) restored 70% of this energy loss.

To examine the effect of the L190P variant in the Na^+^ transport properties of the TaHKT1;5-D transporter, *TaHKT1;5-D* cRNA from Mocho de Espiga Branca or Gladius was introduced in *X. laevis* oocytes. When exposed to different concentrations of Na^+^ glutamate (1 and 30 mM Na^+^), the oocytes with *TaHKT1;5-D* from Gladius had a significantly higher Na^+^ elicited inward current, whereas those with *TaHKT1;5-D* from Mocho de Espiga Branca had limited current, similar to the H_2_O-injected oocytes (Figure 4a). The TaHKT1;5-D from Gladius showed a positive reversal potential shift when exposed to 30 mM Na^+^ which was not observed with TaHKT1;5-D L190P from Mocho de Espiga Branca (Figure 4a).

**Figure 4.**
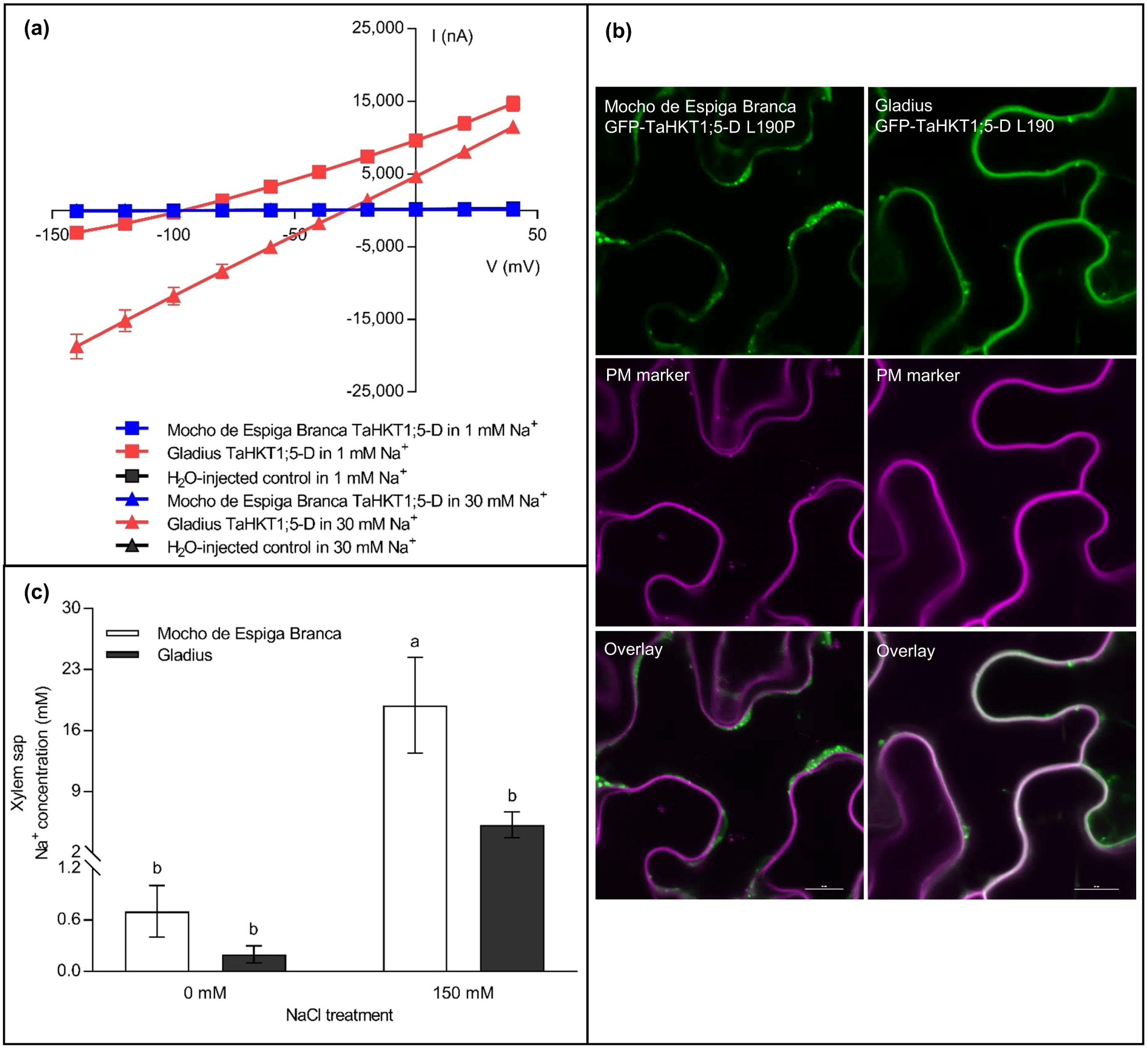
The physiological characterisation of L190P variation in TaHKT1;5-D and the evaluation of the xylem sap Na^+^ concentration in Mocho de Espiga Branca. **(a)** Current-voltage (I-V) curve observed from Mocho de Espiga Branca (blue) or Gladius (red) *TaHKT1;5-D* cRNA-injected or H_2_O-injected (black) *X. laevis* oocytes exposed to 1 mM Na^+^ and 30 mM Na^+^ glutamate at a voltage range from −120 to +40 mV. Data represents means ± SEM (*n* = 3-5). **(b)** Transient co-expression of GFP-TaHKT1;5-D variants with a plasma membrane marker in *N. benthamiana* epidermal cells. Leaves were co-infiltrated with *Agrobacterium tumefaciens* strains harbouring either GFP-TaHKT1;5-D L190 (Gladius) or L190P (Mocho de Espiga Branca) and a plasma membrane marker CBL1n-RFP. GFP signal is shown in green in the left panel while RFP-signal is shown in magenta in the middle panel. Representative images are shown. Scale bars = 10 μm. **(c)** The xylem sap Na^+^ concentration of Mocho de Espiga Branca and Gladius under 0 and 150 mM NaCl concentrations. Xylem sap was collected from hydroponically grown plants 21 days after 0 and 150 mM NaCl was applied at the emergence of 4^th^ leaf. Bars with different letters indicate significant differences determined by two-way ANOVA with Tukey’s multiple comparison test at *p* ≤ 0.05.

Transient expression of N-terminally GFP-tagged TaHKT1;5-D variants in *Nicotiana benthamiana* leaves revealed differences in GFP-signal pattern (Figure 4b). The majority of GFP-TaHKT1;5-D signal co-localised with the plasma membrane (PM) marker CBL1n-RFP, and a minor fraction to mobile subcellular organelles (Figure 4b). GFP-signal in leaves infiltrated with GFP-TaHKT1;5-D L190P, however, localised to internal cell structures, including a faint cytosolic signal and brighter non-mobile structures (Figure 4b). Suggesting the TaHKT1;5-D L190P might get degraded; possibly due to instability of the protein as revealed by homology modelling.

The Na^+^ transport properties of the TaHKT1;5-D L190P variant was further investigated by comparing the xylem sap Na^+^ concentrations of Mocho de Espiga Branca and Gladius grown hydroponically under 0 and 150 mM NaCl. Mocho de Espiga Branca accumulated a 3.5 folds greater xylem sap Na^+^ than Gladius at 150 mM NaCl, while no significant differences were observed at 0 mM NaCl (Figure 4c). There was no significant difference for xylem sap K^+^ and Cl^−^ concentrations for either cultivar (Figure S4a,b).

The ability of Mocho de Espiga Branca to influx or efflux Na^+^ at the root elongation zone was assessed in five day old seedlings using a basal salt medium (BSM) with ion fluxes monitored for 25 min. In the first minute after being exposed to 0.6 mM NaCl, Mocho de Espiga Branca and Gladius had an increase in net Na^+^ efflux up to 10000 and 35000 nmol/m^2^ s^1^ respectively compared to the BSM with 100 mM NaCl (Figure 5a). However, Mocho de Espiga Branca changed within the first minute to Na^+^ influx, while Gladius maintained Na^+^ efflux for the duration of measurement (Figure 5a). The transient net K^+^ efflux was greater in Mocho de Espiga Branca than Gladius and the efflux rates gradually dropped to below 1000 nmol/m^2^ s^1^ in both cultivars (Figure 5b). Mocho de Espiga Branca had a net H^+^ influx for 25 min, while Gladius initially became a net influxer for 8 min and then reverted to being a net effluxer for the remainder of the experiment after exposure to 0.6 mM Na^+^ (Figure 5c). Overall, Mocho de Espiga Branca had greater total Na^+^ influx, K^+^ efflux and H^+^ influx compared to Gladius at the root elongation zone (Figure 5 d,e,f).

**Figure 5.**
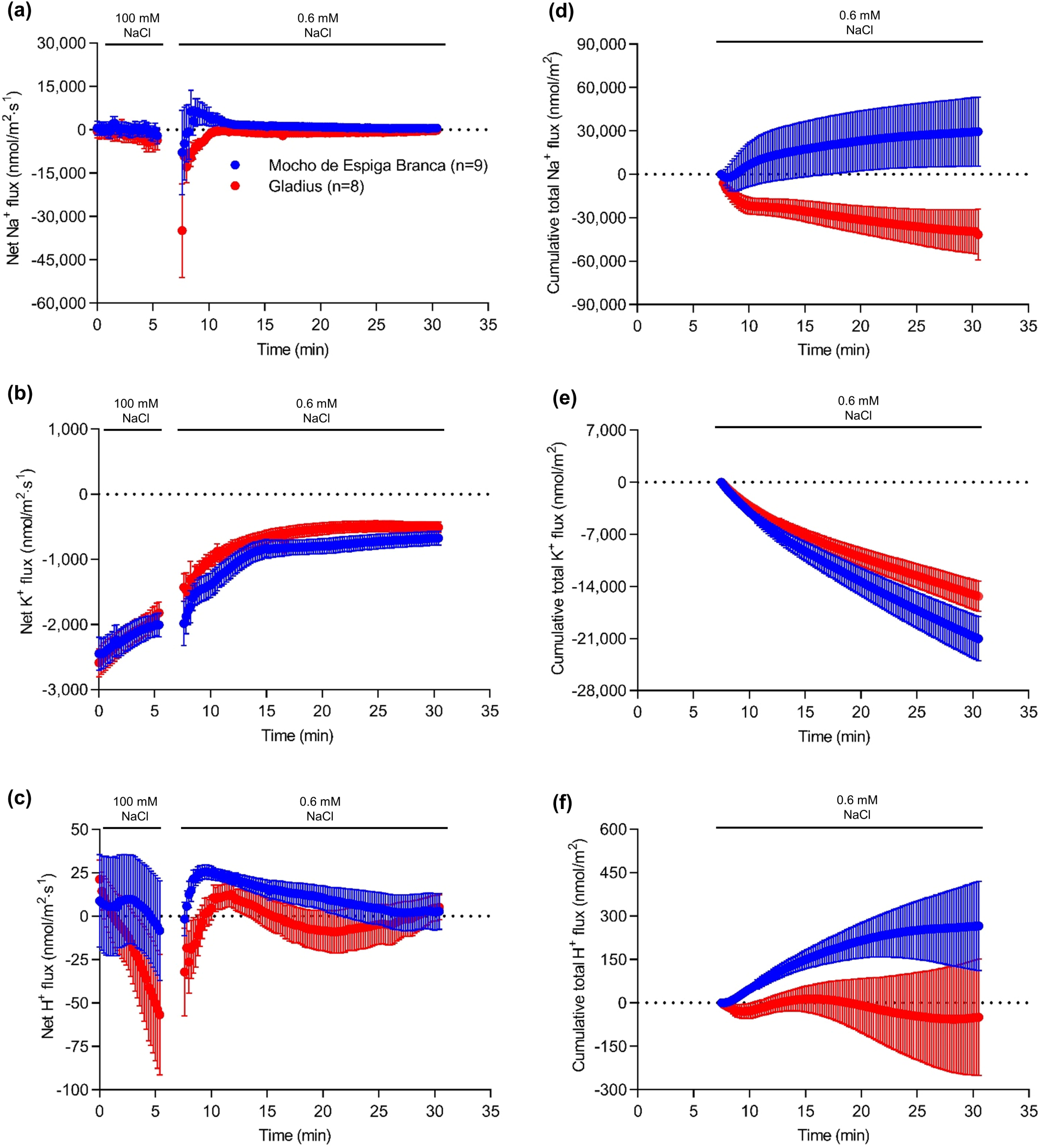
Ion fluxes measured at the root elongation zone after removal from 100 mM NaCl. Six to seven days old Mocho de Espiga Branca or Gladius seedlings were treated with 100 mM NaCl for two days before removal from the solution, the resultant ion fluxes were measured at the elongation zone (between 200 to 600 µm from the root cap) of the primary root of the plants for 25 min. Net **(a)** Na^+^; **(b)** K^+^ and **(c)** H^+^ fluxes. Cumulative total **(d)** Na^+^; **(e)** K^+^ and **(f)** H^+^ fluxes over 25 min. Data presented as means ± SEM (*n* = 8-9).

In contrast, both cultivars had net Na^+^ influx for the first few minutes followed by a gradual net efflux at the maturation zone of the primary root (Figure S5a). As expected salt stress induced K^+^ efflux in both cultivars (Figure S5b). However, there was no significant difference in net or total Na^+^ and K^+^ fluxes observed between the two cultivars, while Mocho de Espiga Branca had 2 folds greater net H^+^ efflux than Gladius (Figure S5 a-f).

## Discussion

A bread wheat landrace, Mocho de Espiga Branca, was identified with significantly higher leaf Na^+^ concentrations and the ability to maintain growth under high salinity. DNA sequencing revealed a naturally occurring SNP in the coding region of the Na^+^ transporter TaHKT1;5-D, resulting in a L190P amino acid residue change. This variation disrupted the capability of TaHKT1;5-D to retrieve Na^+^ from the xylem, due to the protein being unable to transport Na^+^ and/or degradation of the TaHKT1;5-D L190P variant.

3D structural modelling of TaHKT1;5-D and TaHKT1;5-D L190P revealed the important role of the L190P substitution in determining the functional properties of Na^+^ transport in TaHKT1;5-D. The lack of the cooperative binding networks around α-helix 4 of TaHKT1;5-D L190P within the P190 environment (Figure 3d) imposes a severe structural rigidity on the 3D folding of this transporter. The more obtuse packing angle between α-helix 4 and α-helix 5 of TaHKT1;5-D L190P compared to that in TaHKT1;5-D, indicated that the proline position affects packing of α-helices in this specific environment (Figure 3d). Therefore, this α-helix is unlikely to function properly in the structural context preventing Na^+^ ion conductance. In TaHKT1;5-D, a positive correlation was identified between structural characteristics of α-helix 4 and α-helix 5 (trends in angles based on α-helical planes), differences in Gibbs free energies of forward (L190P) and reverse (P190L) mutations and the ability to conduct Na^+^. The inability of the Mocho de Espiga Branca L190P variant to transport Na^+^ was confirmed by expressing the gene in *X. laevis* oocytes (Figure 4a).

Studies also identified differences between the location of the Mocho de Espiga Branca L190P variant and common TaHKT1;5-D. Unlike common TaHKT1;5-Ds, which are localised on the plasma membrane of stelar cells, the L190P variant exhibited greater localisation of GFP signal in internal, non-moving structures (Figure 4b), suggesting the protein is being retained internally and/or is being targeted for degradation. Whether the TaHKT1;5-D L190P is functional and/or not on the correct membrane, Mocho de Espiga Branca will not be able retrieve Na^+^ from the xylem (Figure 4c), explaining why this accession has high leaf blade and leaf sheath Na^+^ (Figure 2b,c).

This work is, to our knowledge, the first that shows a naturally occurring mutation in *TaHKT1;5* directly effects both the Na^+^ transport properties of the protein and the plant phenotype. Previously, differences in the amino acid sequences between Nipponbare and Pokkali *OsHKT1;5* were hypothesised to be responsible for differences in shoot Na^+^ accumulation, however, the transport properties of these proteins were not directly tested (Cotsaftis *et al*., 2012), a similar observation was recently made in barley (van Bezouw, Janssen, Ashrafuzzaman, Ghahramanzadeh, Kilian, Graner, Visser & van der Linden, 2019). Similarly artificially induced mutations in TmHKT1;5-A, which occurred during cloning of the gene, were shown to disrupt Na^+^ transport properties in *X. laevis* but this was not linked to a plant phenotype (Xu *et al*., 2018). A similar natural HKT1;5 variant (L189P) has recently been identified in barley accessions accumulating high grain Na^+^ concentration, which also lacked the ability to transport Na^+^ in *X. laevis* oocytes and was similarly shown to be on internal subcellular structures (Houston et al. unpublished).

Both Mocho de Espiga Branca and Gladius had similar concentrations of K^+^ in the xylem sap (Figure S4a), however Mocho de Espiga Branca accumulated less K^+^ in the leaf blade and sheath (Figure S2a,b). The greater K^+^ efflux at the root elongation zone in Mocho de Espiga Branca (Figure 5b,e) suggests the significant reduction in K^+^ in the leaf blade and sheath is associated with increased K^+^ leakage from the roots. Root Na^+^ and K^+^ concentrations were similar between the two cultivars (Figure 2d and Figure S2c), even though there was a greater root Na^+^ influx (Figure 5a,d), increased shoot Na^+^ (Figure 2b,c), a higher root K^+^ efflux (Figure 5b,e) and reduced shoot K^+^ (Figure S2a,b) in Mocho de Espiga Branca. Bread wheat may have a mechanism, which enables the maintenance of optimal root Na^+^ and K^+^ concentrations contributing to the tolerance of the whole plant to salinity stress. In both cultivars, a high concentration of Cl^−^ was transported in the xylem sap (Figure S4b), which accumulated in the leaf sheath to a greater extent than the leaf blade (Figure S2d,e). This is in agreement with previous findings of Cl^−^ partitioning into the leaf sheath in response to salinity and suggests that the leaf sheath may have an important role in Cl^−^ exclusion preventing it from accumulating to toxic concentrations in the leaf blade (Boursier & Läuchli, 1989, Boursier, Lynch, Lauchli & Epstein, 1987).

Based on the observed functional defects in Na^+^ transport resulting from the TaHKT1;5-D L190P variant and the ion analysis findings in this study, we propose a model to compare root-to-shoot ion transport between Mocho de Espiga Branca and Gladius (Figure 6). Due to the naturally occurring SNP in *TaHKT1;5-D*, we suggest that Mocho de Espiga Branca has impaired retrieval of Na^+^ from the root xylem which results in a greater influx of Na^+^ in the xylem sap and a higher accumulation in the leaf blade and sheath compared to Gladius (Figure 6).

**Figure 6.**
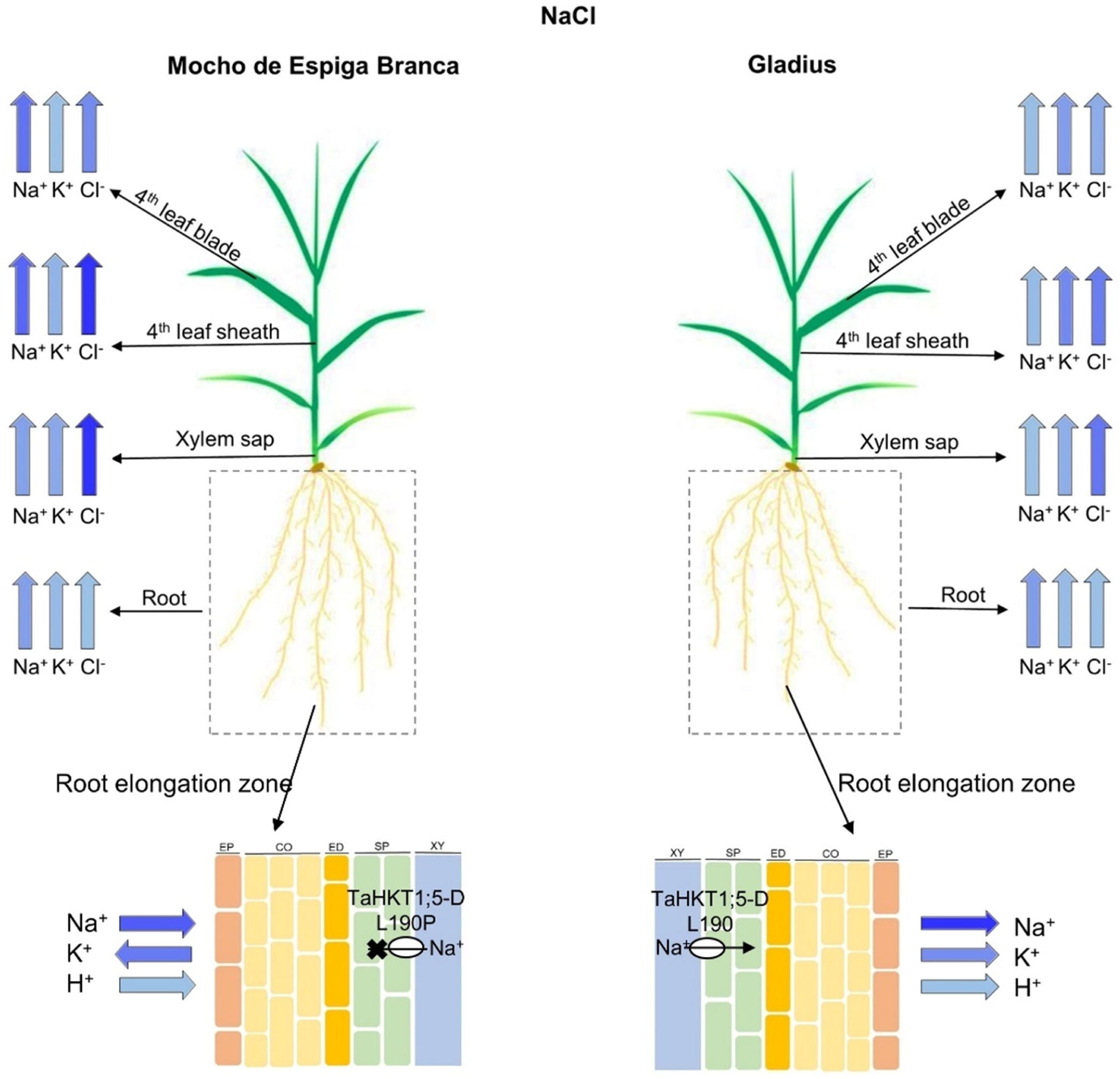
Ion transport model for Mocho de Espiga Branca and Gladius plants under NaCl stress. Colour intensity of the arrows is proportional to the measured ion concentrations, with a greater intensity representing a higher concentration. At the root elongation zone, the direction of the arrow indicates the direction of the ion flux, EP = epidermis, CO = cortex, ED = endodermis, SP = stellar parenchyma and XY = xylem apoplast. TaHKT1;5-D L190P in Mocho de Espiga Branca and TaHKT1;5-D L190 in Gladius are indicated as a white circle. The TaHKT1;5-D L190P variant in Mocho de Espiga Branca leads to reduced retrieval of Na^+^ from the xylem into the roots resulting in a greater influx of Na^+^ in the xylem sap and a higher accumulation of Na^+^ in the leaf blade and sheath compared to Gladius. Mocho de Espiga Branca also has greater Na^+^ influx at the root elongation zone compared to Gladius. There was no difference in root Na^+^ concentration between the two cultivars. The K^+^ concentration was also similar between Mocho de Espiga Branca and Gladius both in the roots and xylem sap, however, Mocho de Espiga Branca had less K^+^ in the leaf blade and sheath compared to Gladius. In both cultivars, a high concentration of Cl^−^ was transported in the xylem sap and accumulated to a high concentration in the leaf sheath compared to leaf blade.

The results of this study confirm the importance of *TaHKT1;5-D* in the Na^+^ exclusion mechanism of bread wheat to limit the levels of Na^+^ that accumulate. It is evident, however, that *TaHKT1;5-D* does not represent the only mechanism responsible for the salinity tolerance of a whole plant, as Mocho de Espiga Branca maintained similar tolerance to Gladius and Scout despite carrying the non-functional *TaHKT1;5-D*. It appears that although *TaHKT1;5-D* has a key role in Na^+^ exclusion, salinity tolerance in bread wheat may not necessarily be entirely related to the plants ability to maintain a low shoot Na^+^ concentration. The lack of a relationship between shoot Na^+^ concentration and salinity tolerance in bread wheat has been observed in other studies (Genc, McDonald & Tester, 2007, Genc *et al*., 2019). This raises the question of how Mocho de Espiga Branca and these other bread wheat accessions maintain growth despite accumulating a high concentration of shoot Na^+^? There must be other tolerance mechanisms, such as tissue tolerance, that are responsible for the plant’s ability to tolerate high shoot Na^+^ concentrations under salinity. Tissue tolerance mechanisms, such as vacuolar Na^+^ compartmentation, the synthesis of compatible solutes and production of enzymes responsible for reactive oxygen species (ROS) metabolism are reported to be essential in plant growth maintenance (Flowers & Colmer, 2008, Munns & Tester, 2008). Specific signalling pathway mechanisms, such as those for ROS, which have been shown to play a role in regulating vasculature Na^+^ concentrations (Mittler, Vanderauwera, Suzuki, Miller, Tognetti, Vandepoele, Gollery, Shulaev & Van Breusegem, 2011, Suzuki, Koussevitzky, Mittler & Miller, 2012), or Ca^2+^ pathways, which regulate gene expression and protein activities (Kudla, Batistič & Hashimoto, 2010, Thoday-Kennedy, Jacobs & Roy, 2015), could also be important for the plant and/or cell’s ability to tolerate high concentrations of shoot Na^+^. Therefore, future studies towards improving salinity tolerance of bread wheat should focus on identifying genetics and physiological mechanisms involved in the plant’s tolerance to high shoot Na^+^ (tissue tolerance) rather than preferentially focusing on Na^+^ exclusion. This will now be easier, knowing it is possible for wheat to survive high shoot Na^+^ concentrations.

In summary, this study identified a bread wheat landrace, Mocho de Espiga Branca, that maintains shoot growth while accumulating very high leaf Na^+^ concentrations under salinity - a novel bread wheat line with tissue tolerance. A naturally occurring SNP variation in the coding region of *TaHKT1;5-D* of Mocho de Espiga Branca results in the amino acid residue substitution L190P in the Na^+^ transporter TaHKT1;5-D, and this single substitution appeared to negatively affect the Na^+^ transport function of the protein, which results in high leaf Na^+^ concentrations. Access to both Mocho de Espiga Branca and the CAPS marker tsl2SALTY-4D for tracking the SNP variation in TaHKT1;5-D will enable plant breeders to develop salinity tolerant bread wheat varieties in the future.

## Acknowledgements

This project was funded by the Grains Research and Development Corporation (GRDC): Project UA00145, UA00151, and the GRDC and the International Wheat Yield Partnership (IWYP): Projects IWYP39/ACP0009; IWYP60/ANU00027. The Australian Centre for Plant Functional Genomics (ACPFG) was jointly funded by the Australian Research Council (ARC) and the GRDC, SW was supported by the ARC DE160100804. The Plant Accelerator^®^, Australian Plant Phenomics Facility, is funded under the National Collaborative Research Infrastructure Strategy (NCRIS). We acknowledge the Cereal Genetic Resource Centre (GRC) INRA Clermont-Ferrand, Assoc. Prof. Kenneth Chalmers and Dr. Melissa Garcia for providing seeds. We thank The Plant Accelerator^®^ for assisting with the glasshouse experiments and Adelaide microscopy for assistance with imaging. We also thank Prof. Rana Munns (CSIRO, Black Mountain, Canberra and University of Western Australia, Perth, Australia) for valuable comments and discussion on the manuscript. CB thanks the China Scholarship Council and the University of Adelaide Joint Postgraduate Scholarships Program for her PhD stipend, and acknowledges the Research Travel Scholarship and the Global Learning Travel Grant at the University of Adelaide and The Plant Nutrition Trust for financial support to attend conferences. The authors declare that they have no conflict of interest.

