## Supporting information for "A single nucleotide substitution in *TaHKT1;5-D* controls shoot Na^+^ accumulation in bread wheat"

**Figure S1.** Plant biomass and salinity tolerance of plants in relation to their 4$^{th}$ leaf $Na^+$ concentration in Mocho de Espiga Branca and two Australian cultivars Gladius and Scout.

**Figure S2.** $K^+$ and $Cl^-$ concentrations in the 4$^{th}$ leaf blade, sheath, and roots of Mocho de Espiga Branca, Gladius and Scout.

**Figure S3.** Expression of *TaHKT1;5-D* in the root tissue and CAPS marker tsl2SALTY-4D genotyping of Mocho de Espiga Branca, Gladius and Scout.

**Figure S4.** Xylem sap $K^+$ and $Cl^-$ concentration of Mocho de Espiga Branca and Gladius.

**Figure S5.** Ion fluxes measured at root mature zone after a sudden exposure to 100 mM NaCl concentration.

**Table S1.** Refer to Excel spreadsheet.

**Table S2.** Genotyping of Mocho de Espiga Branca, Gladius, Scout and 68 bread wheat diversity lines using the CAPS marker tsl2SALTY-4D.

**Table S3.** Primers used for *TaHKT1;5-D* coding sequence amplification and sequencing.

**Supporting Figures:**

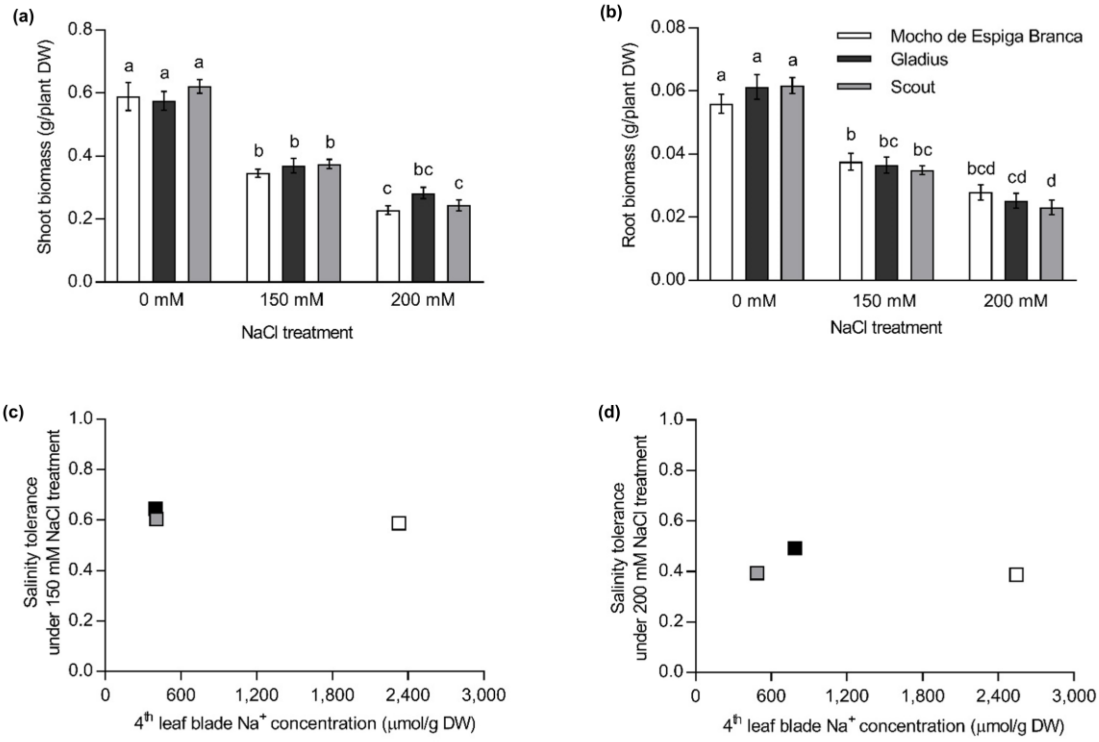


**Figure S1.** Plant biomass and salinity tolerance of plants in relation to their 4$^{th}$ leaf $Na^+$ concentration in Mocho de Espiga Branca and two Australian cultivars Gladius and Scout in a hydroponic system with 0, 150 and 200 mM NaCl concentrations. Plant biomass, salinity tolerance and the 4$^{th}$ leaf $Na^+$ concentration were determined 21 days after 0, 150 and 200 mM NaCl treatments were applied at the emergence of the 4$^{th}$ leaf. Salinity tolerance is defined as the shoot dry weight under 150 or 200 mM NaCl divided by the shoot dry weight under 0 mM NaCl. **(a)** Shoot dry weight (g DW/plant); **(b)** Root dry weight (g DW/plant). **(c)** Salinity tolerance of Mocho de Espiga Branca, Gladius and Scout in relation to their 4$^{th}$ leaf $Na^+$ concentration under 150 and **(d)** 200 mM NaCl. Data are presented as means ± SEM ($n$ = 14). Bars with different letters indicate significant differences determined by two-way ANOVA with Tukey's multiple comparison test at $p \leq 0.05$.

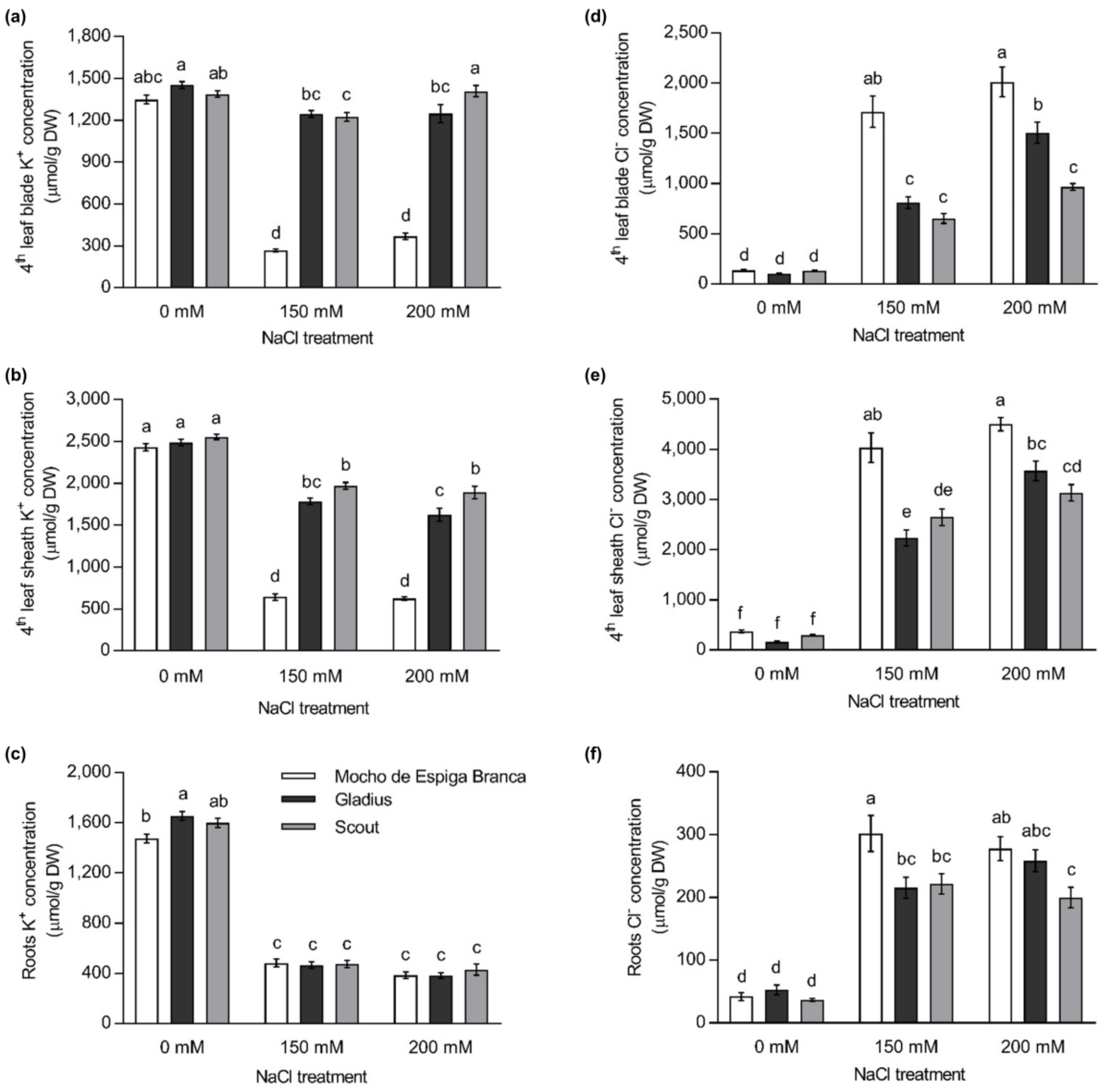


**Figure S2.** $K^+$ and $Cl^-$ concentrations in the 4th leaf blade, sheath, and roots of Mocho de Espiga Branca, Gladius and Scout in a hydroponic system. $K^+$ concentrations in **(a)** 4th leaf blade, **(b)** 4th leaf sheath, **(c)** roots; and $Cl^-$ concentration in **(d)** 4th leaf blade, **(e)** 4th leaf sheath and **(f)** roots determined 21 days after 0, 150 and 200 mM NaCl treatments were applied at the emergence of the 4th leaf. Data are presented as means ± SEM ($n$ = 14). Bars with different letters indicate significant differences determined by two-way ANOVA with Tukey's multiple comparison test at $p \leq 0.05$.

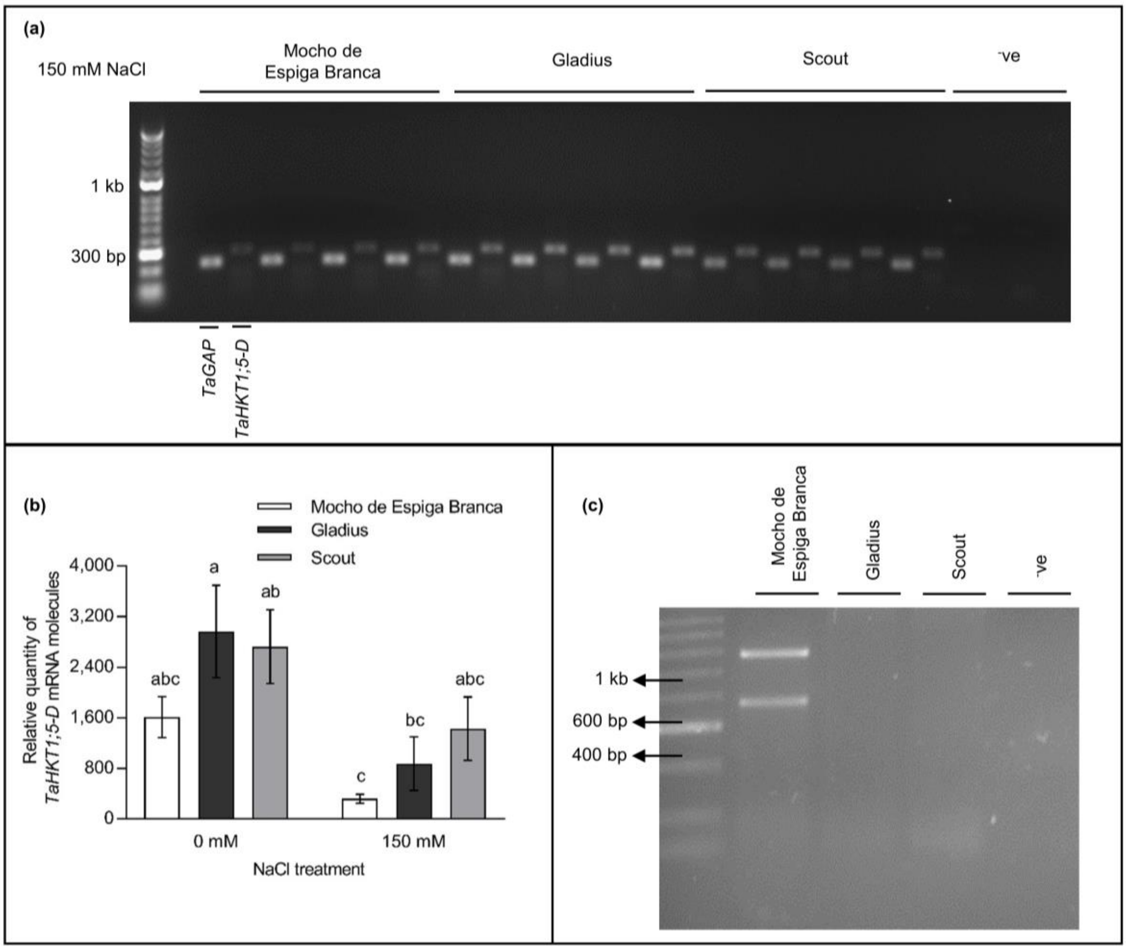


**Figure S3.** Expression of *TaHKT1;5-D* in the root tissue and CAPS marker tsl2SALTY-4D genotyping of Mocho de Espiga Branca, Gladius and Scout. Roots from Mocho de Espiga Branca, Gladius and Scout were hydroponically grown for 21 days under 0 and 150 mM NaCl concentrations which were applied at the emergence of the 4$^{th}$ leaf. **(a)** Expression of *TaHKT1;5-D* in Mocho de Espiga Branca, Gladius and Scout in comparison to positive control (*TaGAP*) and a negative control ($^{-}$ve, Milli-Q water) visualized on 1% (w/v) agarose gel. **(b)** Real-time PCR analysis of *TaHKT1;5-D* mRNA transcript levels relative to *TaActin* and *TaCyclophilin* in root tissue of Mocho de Espiga Branca, Gladius and Scout. Data represents means ± SEM ($n$ = 15). Bars with different letters indicate significant differences determined by two-way ANOVA with Tukey's multiple comparison test at $p \leq 0.05$. **(c)** Agarose gel visualization of the CAPS marker tsl2SALTY-4D genotyping of Mocho de Espiga Branca (T:T), Gladius (C:C) and Scout (C:C). Mocho de Espiga Branca has two fragments at 573 and 372 bp, and Gladius and Scout have a single fragment at 945 bp; $H_2O$ sample is a negative control.

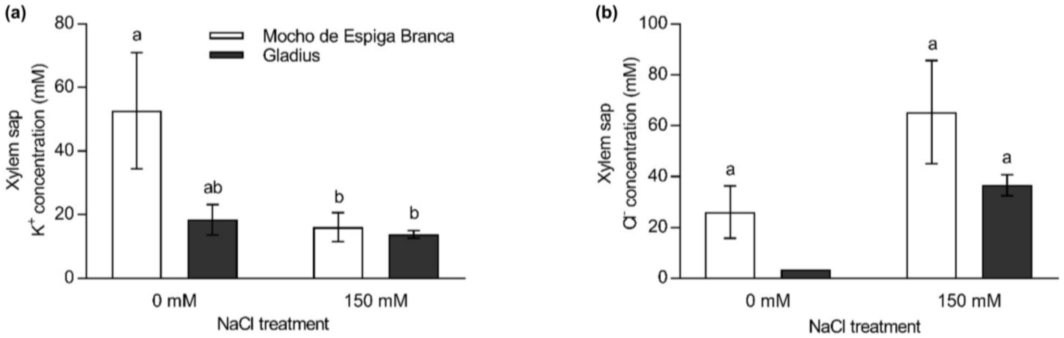


**Figure S4.** Xylem sap $K^+$ and $Cl^-$ concentration of Mocho de Espiga Branca and Gladius under 0 and 150 mM NaCl concentrations. Xylem sap was collected from hydroponically grown plants 21 days after 0 and 150 mM NaCl were applied at the emergence of $4^{th}$ leaf. **(a)** Xylem sap $K^+$ concentration (mM) and **(b)** xylem sap $Cl^-$ concentration (mM). Data presented as mean ± SEM ($n$ = 6-8 except for $Cl^-$ concentration in Gladius under 0 mM NaCl where $n$ = 2). Bars with different letters indicate significant differences determined by two-way ANOVA with Tukey's multiple comparison test at $p \leq 0.05$.

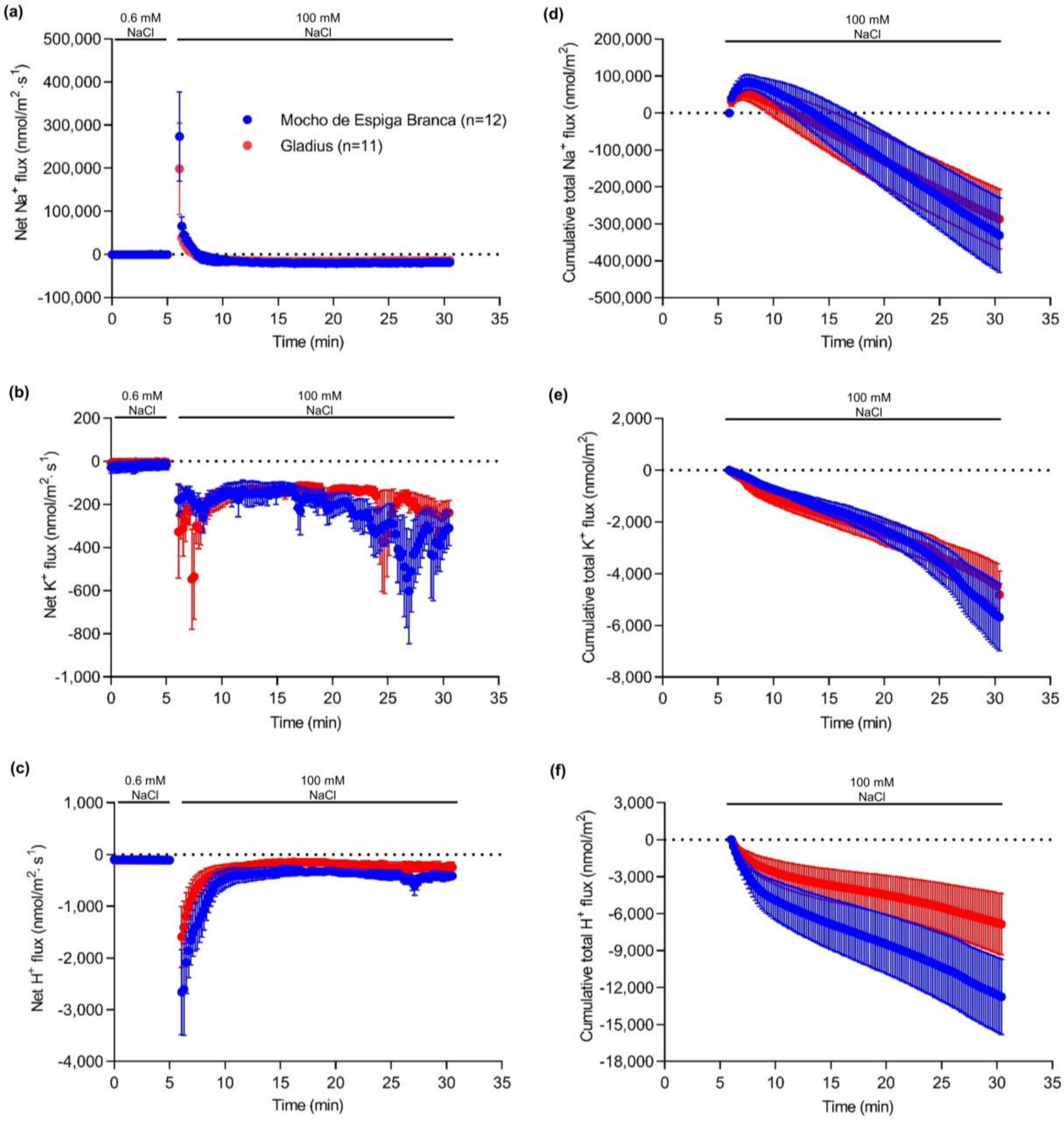


**Figure S5.** Ion fluxes measured at root mature zone after a sudden exposure to 100 mM NaCl concentration. Five-day-old Mocho de Espiga Branca or Gladius seedlings were treated with 100 mM NaCl, and the resultant ion fluxes were measured at the root maturation zone (approximately two cm above the root apex) of the primary root for 25 min. Net **(a)**, $Na^+$; **(b)**, $K^+$ and **(c)**, $H^+$ fluxes. Cumulative total **(d)**, $Na^+$; **(e)**, $K^+$ and **(f)**, $H^+$ fluxes over 25 min. Data are presented as means ± SEM (*n* = 11-12).

**Supporting Tables:**

**Table S1.** Screening of 75 bread wheat lines for salinity tolerance. (Refer to Excel spreadsheet).

**Table S2:** Genotyping of Mocho de Espiga Branca, Gladius, Scout and 68 bread wheat diversity lines for the SNP (T/C) resulting in TaHKT1;5-D L190P variation using the CAPS marker tsl2SALTY-4D designed for the SNP.

| ***TaHKT1;5-D*** **SNP allele** | **Bread wheat accessions** |
|---|---|
| C:C | Mocho de Espiga Branca |
| T:T | Gladius, Scout, Odessa EXP STA 17413, Pitic 62, W 98, Buck Atlantico, LW 337, G 72300, LW 333, Siete Cerros, N 46, Ciano 79, Meira, Menflo, Sokool, Persia 21, LW 335, H 1534, M708/G25/N163, Odessa EXP STA 19565, LW 336, Coppadra, S975-A4-A1, Frontana 3671, Palestinskaya, Taferstat, BuckBuck 'S', India 231, Persia 7, Suzuki 24, Granarolo, India 227, India 322, Suzuki 17, Persia 4, Zilve, Vorobey, 6/01/2003, Academie de Pekin, H 1685, Bahatane 87, Optata 85, Jupateco 73, H 1440, N67M2, Yecora 70, Thori 212-var 8/1, Estanzuela Dorado, LW 349, Cul 10, Asia Minor 71, Inia 66, Suzuki 23, Suzuki 12, LW 334, Suzuki 3, LW 369, EMU'S', PBW 343, Genaro F81, Pamukale, Daeraad, BT 2281, Cul 9, BT 2277, 117-var 12/564, Andes 56, Suzuki 9, Tacupeto 2001, Roelfs F2007 |

**Table S3.** Primers used for *TaHKT1;5-D* coding sequence amplification and sequencing. FP = Forward Primer; RP = Reverse Primer; Primers marked '*' are from Byrt, et al (2014).

| **Primers** | **Start (bp)** | **Length (bp)** | **Sequences (5'- 3')** |
|---|---|---|---|
| *cTaHKT1,5-D_FP_1* | 1 | 22 | ATGGGTTCTTTGCATGTCTCCT |
| *cTaHKT1,5-D_RP_328* | 328 | 20 | GCATGCTCGTGAACACCTCG |
| *cTaHKT1;5-D_FP_429* | 429 | 20 | CCTAGAGCTCGCTGTTACCA |
| *cTaHKT1; 5-D_FP_693** | 693 | 17 | CTGCGGCTTCGTCCCGA |
| *cTaHKT1; 5-D_FP_1194** | 1194 | 24 | CGACCAGAAAAGGATAACAAGCAT |
| *cTaHKT1; 5-D_RP_1551* | 1551 | 23 | TTATACTATCCTCCATGCCTCGC |
